# Recurrent emergence of carbapenem resistance in *Klebsiella pneumoniae* mediated by an inhibitory *ompK36* mRNA secondary structure

**DOI:** 10.1101/2022.01.05.475072

**Authors:** Joshua L. C. Wong, Sophia David, Julia Sanchez-Garrido, Jia Z. Woo, Wen Wen Low, Fabio Morecchiato, Tommaso Giani, Gian Maria Rossolini, Stephen J. Brett, Abigail Clements, David M Aanensen, Silvi Rouskin, Gad Frankel

## Abstract

Outer membrane porins in Gram-negative bacteria facilitate antibiotic influx. In *Klebsiella pneumoniae* (KP), modifications in the porin OmpK36 are implicated in increasing resistance to carbapenems. Analysis of large KP genome collections, encompassing major healthcare-associated clones, revealed the recurrent emergence of a synonymous cytosine to thymine transition at position 25 (25c>t) in *ompK36.* We show that the 25c>t transition increases carbapenem resistance through depletion of OmpK36 from the outer membrane. The mutation attenuates KP in a murine pneumonia model, which accounts for its limited clonal expansion observed by phylogenetic analysis. However, in the context of carbapenem treatment, the 25c>t transition tips the balance towards treatment failure, thus accounting for its recurrent emergence. Mechanistically, the 25c>t transition mediates an intramolecular mRNA interaction between a uracil encoded by 25t and the first adenine within the Shine-Dalgarno sequence. This specific interaction leads to the formation of an RNA stem structure, which obscures the ribosomal binding site thus disrupting translation. While mutations reducing OmpK36 expression via transcriptional silencing are known, we uniquely demonstrate the repeated selection of a synonymous *ompK36* mutation mediating translational suppression in response to antibiotic pressure.

## Introduction

Classical outer membrane (OM) porins in Gram-negative bacteria enable non-specific bidirectional diffusion between the periplasm and extracellular environment (Achouak et al., 2001). This has been exploited in antibacterial chemotherapy as porins act as the key entry point for clinically important classes of antibiotics across the otherwise impermeable OM. However, porin modifications that restrict antibiotic entry have evolved in response to this selective pressure (Vergalli et al., 2020), contributing to the rising global burden of resistant bacterial infections, especially among species of *Enterobacteriaceae* (Bajaj et al., 2016; Dé et al., 2001; Wong et al., 2019; Fajardo-Lubián et al., 2019; Tran et al., 2010).

*Klebsiella pneumoniae* (KP) is one of the most clinically significant members of the *Enterobacteriaceae* family and a leading cause of healthcare-associated infections worldwide (Cassini et al., 2019; Vincent et al., 2020). The majority of resistant KP infections are caused by “high-risk” clonal lineages, including sequence types (ST) 258 and 512, which form a dominant clone associated with the KP carbapenemase (KPC) gene (David et al., 2019). Together with plasmid-encoded carbapenemases, modifications to the major OM porins OmpK35 and OmpK36 play a critical role in mediating KP resistance to carbapenems, a class of antibiotics that is vital for the treatment of severe infections. Resistance-associated modifications broadly fall into those that structurally alter the porin channel and those that abolish or reduce OmpK36 expression.

Structural alterations in OmpK36 are mediated by amino acid insertions into a region of the porin called loop 3 (L3). These insertions narrow the luminal diameter and restrict substrate diffusion, including antibiotics (Wong et al., 2019; Fajardo-Lubián et al., 2019).

L3 insertions are relatively prevalent among clinical KP isolates, having been found among 12.3% (192/1557) of isolates from a diverse public genome collection (Fajardo-Lubián et al., 2019). We previously showed that the most common L3 insertion, a di-amino acid insertion (Glycine-Aspartate, GD), results in a 16-fold increase in minimum inhibitory concentration (MIC) to meropenem (Wong et al., 2019).

In addition to structural modifications, carbapenem resistance is also achieved by absent or reduced OmpK36 expression. Absent expression, which can be achieved by gene truncation, results in high levels of resistance but comes at a significant *in vivo* fitness cost (Wong et al., 2019; Fajardo-Lubián et al., 2019; Tsai et al., 2011). Reduced OmpK36 expression has been reported to occur by multiple mechanisms that all result from transcriptional silencing, including *ompK36* promoter disruption by insertion sequence (IS) elements (Clancy et al., 2013), loss-of-function mutations in *kvrA* (a transcriptional repressor that controls capsule production) (Dulyayangkul et al., 2020) and mutations in *hfq* (a regulatory RNA binding protein) (Chiang et al., 2011). However, the effects on virulence resulting from reduced OmpK36 expression are poorly understood. Moreover, the prevalence and clinical significance of these mechanisms remain unknown as the mutations identified to date were either restricted to clinical isolates from a single centre (promoter insertion) or evolved during *in vitro* selection or genetic deletion experiments (*kvrA* and *hfq*).

Here we identify and describe a carbapenem resistance mechanism that has been repeatedly employed by clinical KP isolates to post-transcriptionally alter OmpK36 abundance via synonymous mutations in the open reading frame (ORF). Our study spans the identification of one such key mutation (25c>t) in *ompK36* through large-scale bioinformatic approaches, the assessment of its effects on carbapenem susceptibility and virulence using murine pneumonia models, and finally the determination of the precise molecular mechanism linking synonymous SNPs with protein abundance. In particular, we show that the 25c>t mutation results in the formation of a stem structure in the *ompK36* mRNA that obscures the Shine Dalgarno sequence (SDS), reduces OmpK36 translation and abundance, and increases carbapenem resistance as a consequence. This work provides the first example of the functional use of these inhibitory mRNA structures in regulating protein expression during bacterial adaptation, and characterises a novel, clinically-important resistance mechanism in KP.

## Results

### Recurrent emergence of a synonymous 25c>t ompK36 mutation

We curated and analysed a collection of 1450 public KP genomes belonging to the major healthcare-associated clone composed of STs 258, 512 and other closely related derivatives (https://microreact.org/project/1vWbaqARPRNc55n4yfdLyQ-ompk36; **Table S1**). The collection comprises isolates gathered between 2003 and 2018, largely in the Americas, Europe and the Middle East, where rampant spread of ST258/512 in healthcare institutions has been reported (Adler et al., 2017; Giakkoupi et al., 2011; Giani et al., 2013; Kitchel et al., 2009; Rojas et al., 2017). We unambiguously identified the *ompK36* gene in 98.1% (1422/1450) of the genomes. Among these, the gene was intact in 99.3% (1412/1422). Using the intact *ompK36* sequences, we inferred how each position in this gene has evolved across the ST258/512 population. First, we constructed a phylogeny of the collection using all vertically inherited single nucleotide polymorphisms (SNPs) from a core genome alignment, with recombined regions excluded, to provide an accurate representation of the ancestral relationships between isolates (Figure 1A). We then mapped the variation in *ompK36* onto this phylogeny and predicted the ancestral states of each position in the gene across the tree using a maximum parsimony approach. While 83.4% of nucleotide positions are entirely conserved across *ompK36*, we found two regions with an increased number of base changes (Figure 1B). As anticipated, the largest number of changes occurred in the loop 3 (L3) region comprising amino acid insertions (GD, TD, D and N) (24 occurrences) as well as their reversions (i.e. deletions) (25 occurrences). However, further to this, we observed ten base changes at position 25, all consisting of a synonymous c>t transition (<u>C</u>TG (leucine)-><u>T</u>TG (leucine)). Despite this mutation maintaining the identical amino-acid translation in the Sec-dependent signal sequence (Figure 2A&B), its high frequency of emergence suggests that it has undergone positive selection. The majority of isolates with the 25c>t mutation are sporadically distributed across the core genome phylogeny and occur as singletons or clusters of only two isolates (11/18) (Figure 1A). The remaining seven isolates, collected between 2009 and 2013 from multiple healthcare institutions in the USA, form a monophyletic cluster, indicative of a clonal expansion. We noted that these seven isolates encode an additional synonymous 24c>t mutation in the preceding codon ((CT<u>C</u> (leucine)->CT<u>T</u> (leucine)) (Figure 2B).

**Figure. 1.**
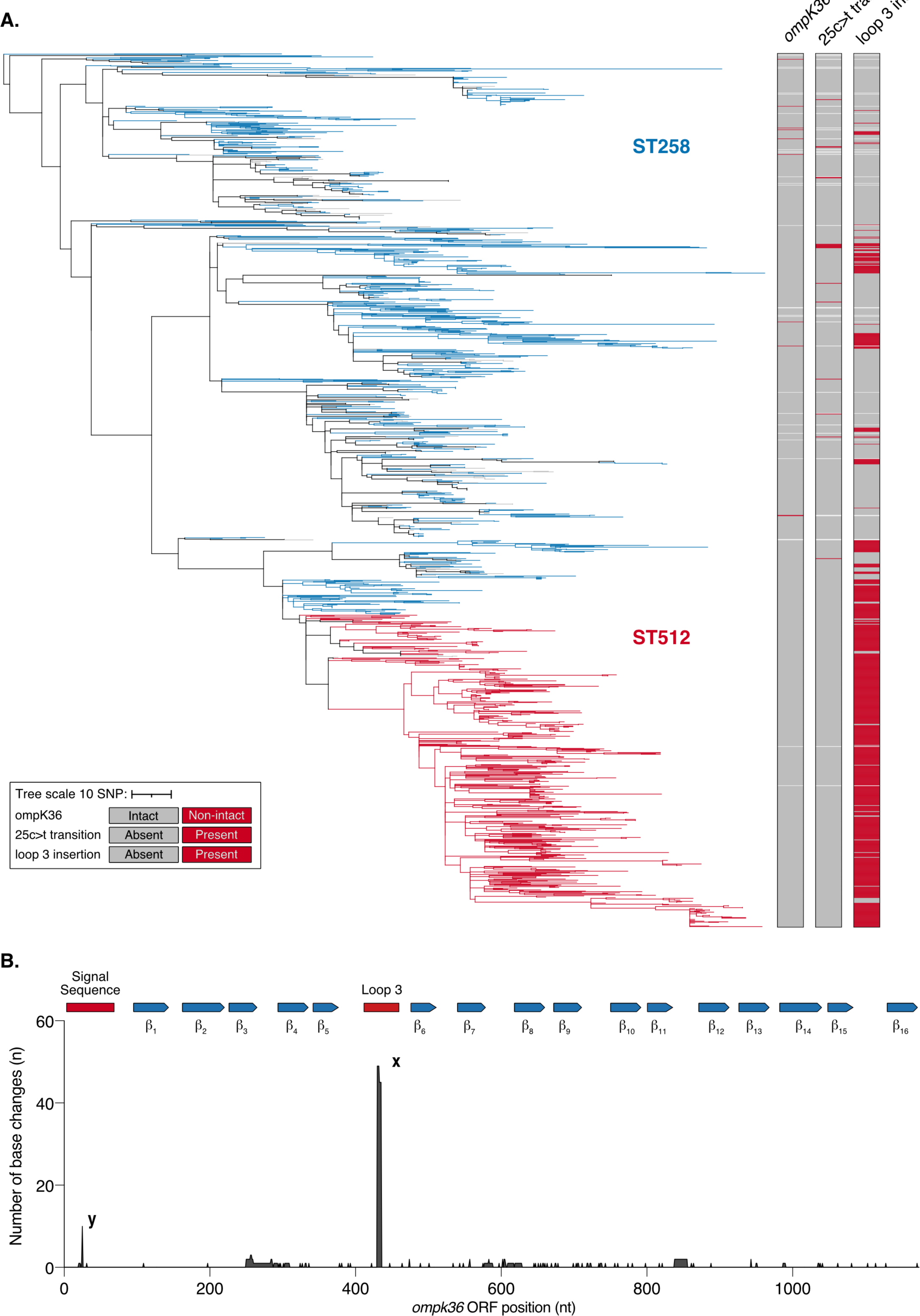
Frequency and phylogenetic distribution of *ompK36* variants in a curated collection of ST258/512 genomes. **A.** Phylogenetic tree of 1450 KP ST258/512 isolates constructed using vertically-inherited single nucleotide polymorphisms (SNPs). Branches are coloured blue if all descendent taxa belong to ST258, red if they belong to ST512, grey if they belong to other closely related derivatives of these STs, or black if they belong to one or more of these different categories. Data columns (L-R) show *ompK36* (intact/non-intact)*, ompK36* 25c>t transition (present/absent) and *ompK36* loop 3 (L3) insertion (present/absent) in each genome. Isolates are marked white if *ompK36* could not be unambiguously identified in the genome (columns 1-3) or if *ompK36* was non-intact (columns 2/3). The scale represents the number of SNPs per variable site. A similar visualisation is available at: https://microreact.org/project/1vWbaqARPRNc55n4yfdLyQ-ompk36#wksn-figure-1a-wong-et-al-2022. **B.** The number of changes (including SNPs, insertions and deletions) at each position in *ompK36* detected across the ST258/512 tree. The protein structure motifs (signal sequence, beta strands 1-16 and L3) are plotted on top of the sequence as a reference. The labelled peaks represent insertions/reversions within the L3 (x) and the 25c>t mutation found within the signal sequence (y).

**Figure. 2.**
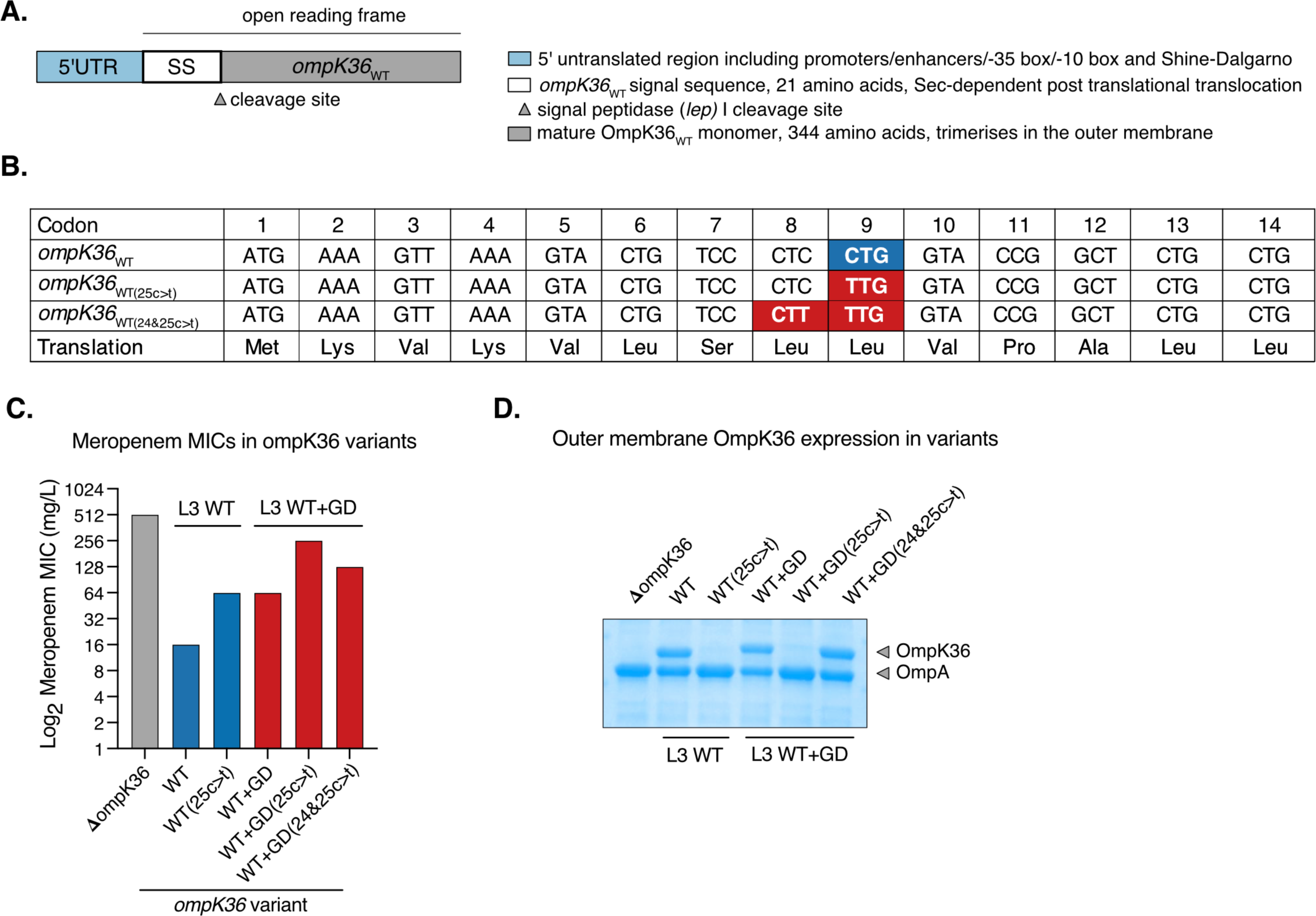
*ompK36*_WT(25c>t)_ leads to increased meropenem MIC and reduced OmpK36 abundance. **A.** Schematic of the *ompK36* locus. **B.** The synonymous 25c>t mutation occurs at codon position 9 (Leu9) resulting in a change of codon from CTG (*ompK36*_WT_) to TTG (*ompK36*_WT(25c>t)_). The 24c>t mutation occurs in position 8 (Leu8) resulting in a synonymous CTC to CTT codon switch. **C.** The 25c>t transition increases the meropenem resistance achieved on both a WT and WT+GD *ompK36* background. The additional 24c>t (WT+GD(24&25c>t)) mutation partially reverses the resistance achieved in the 25c>t transition. All strains harbour a pKpQIL-like plasmid expressing KPC-2 and have *ompK35* deleted. Resistance values represent broth MICs (median of 3 repeats). **D.** Representative Coomassie gel image of outer membrane (OM) preparations demonstrating the reduced OmpK36 expression conferred by 25c>t in both WT and WT+GD backgrounds. The additional 24c>t in WT+GD(24&25c>t) reverses the OM depletion of OmpK36 in WT+GD(25c>t).

**Table 1.** ompK36 variants and KP strains used in this study.

| *ompK36* variants | Nucleotide at | Nucleotide at | Loop 3 (L3) insertion |
| --- | --- | --- | --- |
| *ompK36*<sub>WT</sub> | c | c | Absent |
| *ompK36*<sub>WT(25c>t)</sub> | c | t | Absent |
| *ompK36*<sub>WT+GD</sub> | c | c | Glycine-Aspartate (GD) |
| *ompK36*<sub>WT+GD(25c>t)</sub> | c | t | Glycine-Aspartate (GD) |
| *ompK36*<sub>WT+GD(24&25c>t)</sub> | t | t | Glycine-Aspartate (GD) |
| Strains | *ompK35* | *ompK36* | Carbapenemase |
| KPΔ36 | Δ | Δ | +/- KPC-2* |
| KP36<sub>WT</sub> (+/- KPC-2) | Δ | *ompK36*<sub>WT</sub> | +/- KPC-2* |
| KP36<sub>WT(25c>t)</sub> (+/- KPC-2) | Δ | *ompK36*<sub>WT(25c>t)</sub> | +/- KPC-2* |

\*KPC-2 on pKpQIL-like plasmid

We next searched for the 25c>t mutation in a large, geographically diverse collection of 16,086 KP genomes (https://pathogen.watch/genomes/all?genusId=570&speciesId=573; Table S2) to establish its wider prevalence among sequenced isolates and presence in other (i.e. non-ST258/512) clonal lineages. Among the 14,888 isolates encoding an intact *ompK36* gene, we identified the 25c>t mutation in 2.5% (376/14,888), which includes isolates from 39 STs and 25 countries. Over half (51.3%; 193/376) belong to ST258 while many also belong to other globally-important multi-drug resistant clones (e.g. ST15, 19.9%; ST13, 4.5%; ST11, 2.7%). We observed the 24c>t mutation in 1.1% (161/14,888) of genomes from this collection, and only ever in combination with the 25c>t mutation. Moreover, we also noted that 93.2% (150/161) of *ompK36* sequences containing the 24&25c>t mutations had L3 insertions compared to only 1.9% (4/215) of those with 25c>t only.

### The 25c>t mutation increases meropenem resistance

We modelled the effect of the 25c>t mutation (*ompK36*_WT(25c>t)_) on meropenem resistance by introducing the mutation into *ompK36*_WT_ of our laboratory KP strain ICC8001 (Wong et al., 2019) (see Table 1 for list of *ompK36* variants and strains used in this study). We additionally deleted *ompK35*, as this porin gene is truncated in 99.8% of the ST258/512 genomes, and introduced the KPC-2 carbapenemase (on a pKpQIL-like plasmid) given also the strong association of KPC genes with the lineage (Bowers et al., 2015). We found that expression of *ompK36*_WT(25c>t)_ results in a meropenem minimum inhibitory concentration (MIC) of 64mg/L, 8-fold higher than the resistance breakpoint of 8mg/L (EUCAST, 2021) (Figure 2C). This MIC is also four-fold higher than that seen in KP expressing *ompK36*_WT_ (16mg/L), albeit lower than the very high resistance observed when *ompK36* was deleted (Δ*ompK36,* 512mg/L), indicating that the 25c>t is a carbapenem resistance mutation.

We also investigated the level of resistance conferred by the combination of the 24c>t and 25c>t mutations. While ICC8001 did not tolerate the genomic insertion of the double 24&25c>t mutations on a WT *ompK36* background, we found that they were tolerated when introduced in combination with a L3 GD pore-constricting insertion (*ompK36*_WT_+GD(24&25c>t)). While we could not explain the inability to achieve *ompK36*_WT_(24&25c>t) introduction, it nevertheless reflects the strong association between the combined 24&25c>t mutations and L3 insertions observed in our genomic analysis. Broth microdilution testing of the strain with *ompK36*_WT_+GD(24&25c>t), together with strains encoding *ompK36*_WT_+GD and *ompK36*_WT_+GD(25c>t), revealed that the 25c>t mutation increased the meropenem MIC of the L3 GD insertion strain from 64mg/L (WT+GD) to 256mg/L (WT+GD(25c>t)) and that the double 24&25c>t mutations partially reversed this phenotype, with an MIC of 128mg/L (WT+GD(24&25c>t)) (Figure 2C).

### ompK36*WT(25c>t)* attenuates virulence but tips the balance towards antibiotic therapy failure

The general absence of clonal expansion among isolates with the single 25c>t substitution, despite the frequent emergence of this mutation, suggests that it may have a fitness cost that impedes onward transmission. We tested this hypothesis *in vivo* by infecting mice with 250 CFU of KP encoding either *ompK36*_WT_ (KP36WT) or *ompK36*_WT(25c>t)_ (KP36WT(25c>t)) (Table 1). Inoculation with a strain encoding an *ompK36* deletion (KPΔ36) and PBS were used as controls (Figure 3A). At 48 h post infection, KP36WT, but not KP36WT(25c>t), induced significant weight loss compared to uninfected (PBS) mice or mice infected with KPΔ36, Figure 3B). Bacterial counts in the lungs showed a trend to higher burdens in mice infected with KP36WT than with KP36WT(25c>t), which reached significance in the blood (Figure 3C&D). KPΔ36 demonstrated defects in bacterial survival and proliferation in the host, with significantly lower and/or absent burdens in the lungs and blood (Figure 3C&D). Interferon gamma (IFNγ) is a key cytokine in the defense against KP and measurement of its levels in serum provides a readout of acute inflammatory responses (Moore et al. 2002); following infection with KP36WT, IFNγ responses were significantly higher compared to infection with KP36WT(25c>t) (Figure 3E). Taken altogether, these findings show that the 25c>t transition attenuates KP *in vivo* with reduced bacteraemia and the resulting induction of a diminished host pro-inflammatory immune response.

**Figure. 3.**
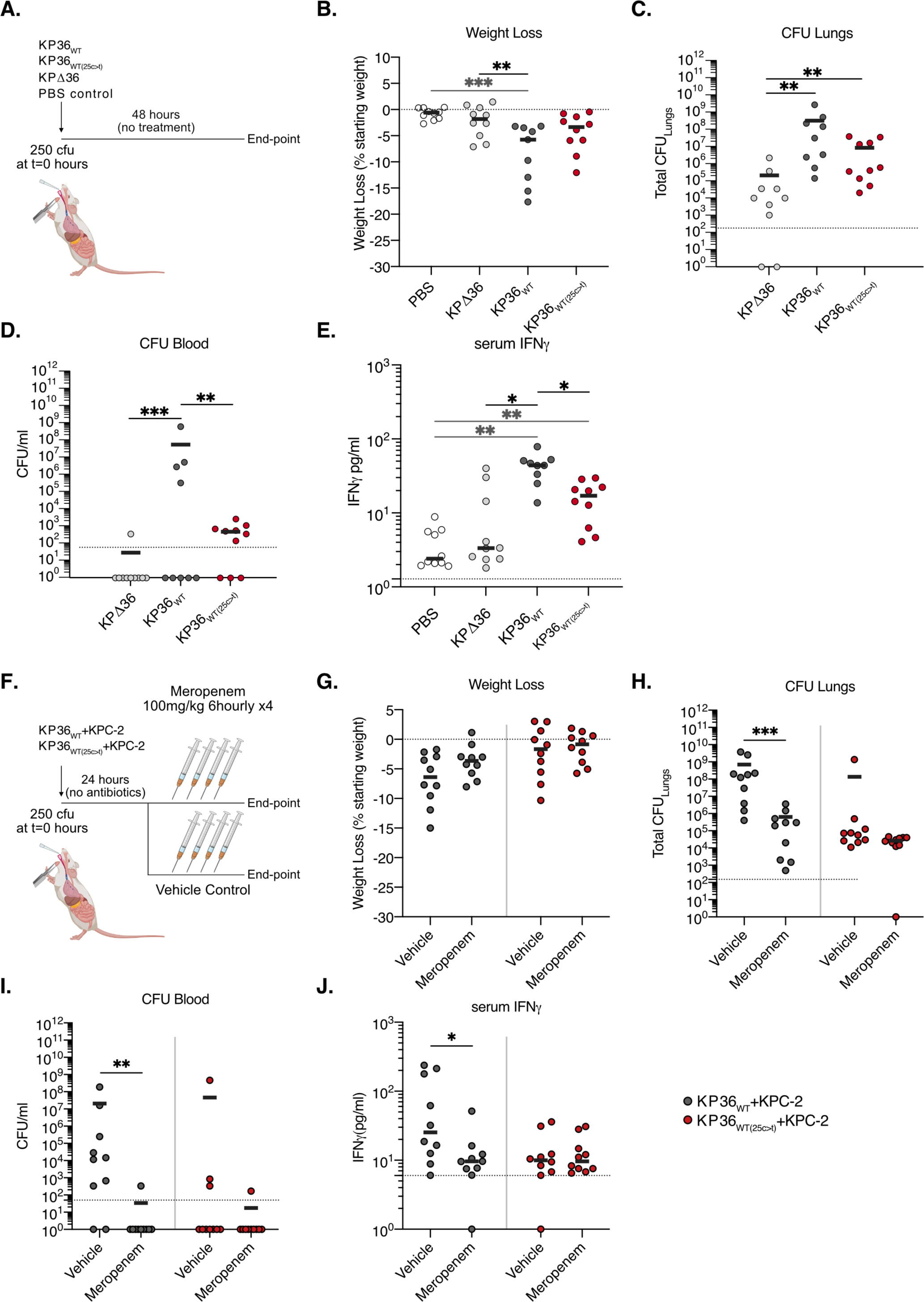
The 25c>t mutation has a fitness cost *in vivo* but is advantageous in the context of meropenem therapy. **A-E.** Mice were infected by intubation and administration of 250 CFU of KP36WT, KP36WT(25c>t) or KPΔ36 strains; mock infection with PBS was used as a control. A schematic of the infection protocol is outlined in panel a. **B.** Animals infected with KP36WT show greater weight loss after 48 h compared to mock-infected animals and those infected with KPΔ36. Weight loss in animals infected with KP36WT(25c>t) is not significantly different to that of the PBS or KPΔ36 groups. **C** and **D**. Enumeration of CFU in lungs and blood, collected at 48 h post infection, reveals that infection with KPΔ36 does not result in high lung bacterial burdens and bacteraemia. Infection with KP36WT results in high pulmonary bacterial burdens (**C**) and bacteraemia (**D**), while infection with KP36WT(25c>t) results in a trend towards lower pulmonary bacterial counts and reduced levels of bacteria in the blood. Dotted line represents the limit of detection of the assay. **E.** Infection with KP36WT or KP36WT(25c>t) leads to increased serum IFNγ compared to mock-infected (PBS) animals. KP36WT(25c>t) infection results in significantly reduced serum IFNγ levels compared to KP36WT infection. **F-J.** Mice were administered with KP36WT+KPC-2 or KP36WT(25c>t)+KPC-2 strains and subjected to a meropenem dosing regimen to assess the success of antibiotic therapy in KP-induced severe pneumonia. A schematic of the infection protocol is outlined in panel **F**. **G.** No changes were observed in weight loss between meropenem and vehicle control therapy when experimental pneumonia induced by either KP36WT+KPC-2 or KP36WT(25c>t)+KPC-2. **H** and **I.** CFU enumeration in lungs (**H**) and blood (**I**) shows significant reduction in meropenem-treated animals infected with KP36WT+KPC-2 compared to vehicle treated animals; levels of bacteraemia were undetectable in all but one meropenem treated animal. Meropenem treatment has no impact on lung or blood CFUs in animals infected with KP36WT(25c>t)+KPC-2. **J.** Serum levels of IFNγ are significantly reduced after meropenem treatment in animals infected with KP36WT+KPC-2; cytokine levels are not affected by meropenem treatment in animals infected with KP36WT(25c>t)+KPC-2. Graphs show median values of 2 biological repeats (4-5 mice per group). Statistical significance was determined by one-way ANOVA with Tukeyśs multiple comparison post-test. *, P < 0.05; **, P < 0.01; ***P < 0.001; ****, P<0.0001 in **B-E**. The statistical significance of the comparisons between vehicle and meropenem-treated animals was determined via a non-parametric Mann-Whitney test. *, P < 0.05; **, P < 0.01; ***P < 0.001 in **G-J**.

We next evaluated if *ompK36*_WT(25c>t)_ provides an advantage compared with *ompK36*_WT_ in the context of meropenem therapy, which would explain the recurrent emergence of this mutation despite its associated fitness cost. We infected mice with either KP36WT or KP36WT(25c>t), both expressing the KPC-2 carbapenemase from a pKpQIL-like plasmid. At 24 h mice either received meropenem (100mg/kg dose) or vehicle control (water) by intraperitoneal injection at six hourly intervals for 24 h (Figure 3F). The experiment was stopped 3 h after the last injection. No significant differences in body weight were observed following infection by either strain with or without meropenem therapy (Figure 3G), consistent with fluid resuscitation provided by meropenem or vehicle administration. However, significant reductions in the bacterial burdens in the lungs and blood were seen following meropenem treatment of KP36WT+KPC-2 but not KP36WT(25c>t)+KPC-2 infection (Figure 3H&I). Furthermore, while the level of serum IFNγ was significantly decreased when KP36WT+KPC-2 infection was treated with meropenem (Figure 3J), no significant differences were observed between antibiotic or mock treated KP36WT(25c>t)+KPC-2 infected mice. These *in vivo* results show whilst the 25c>t mutation in *ompK36* attenuates KP, it provides a selective advantage during carbapenem therapy.

### The 25c>t mutation reduces OmpK36 abundance

We next aimed to understand how the 25c>t mutation mediates an increase in meropenem resistance and an associated decrease in virulence. Since the mutation results in both carbapenem resistance and virulence phenotypes between those observed in normal (*ompK36*_WT_) and absent (Δ*ompK36*) expression, we hypothesized that these phenotypic changes were due to reduced OmpK36 abundance in the OM. Indeed, analysis of isolated OM fractions by SDS-PAGE and Coomassie staining revealed that the strain encoding *ompK36*_WT(25c>t)_ exhibited a substantially reduced abundance of OmpK36 compared to that with *ompK36*_WT_ (Figure 2D), validating this hypothesis. In line with the meropenem MIC results, we also found that this pattern is reversed in the strain encoding *ompK36*_WT_+GD(24&25c>t), which exhibits OmpK36 levels similar to the strain encoding *ompK36*_WT_ (Figure 2D).

### Reduced OmpK36 abundance via the 25c>t mutation occurs independently of codon bias

We sought to describe the precise molecular mechanism linking the synonymous 25c>t mutation to decreased OmpK36 abundance. We first explored whether reduced OmpK36 abundance is due to a decreased translation rate, resulting from a change in codon from the commonly-used CTG to the rarely-used TTG at amino acid position 9 (Leu9). Overall, 64.9% of leucine residues in the ICC8001 genome are encoded by CTG and 5.9% encoded by TTG (Figure 4A); inspection of all the ST258/512 genomes confirmed a similar bias (CTG range: 57.0-65.1%; TTG range: 5.9-9.3%).

**Figure 4.**
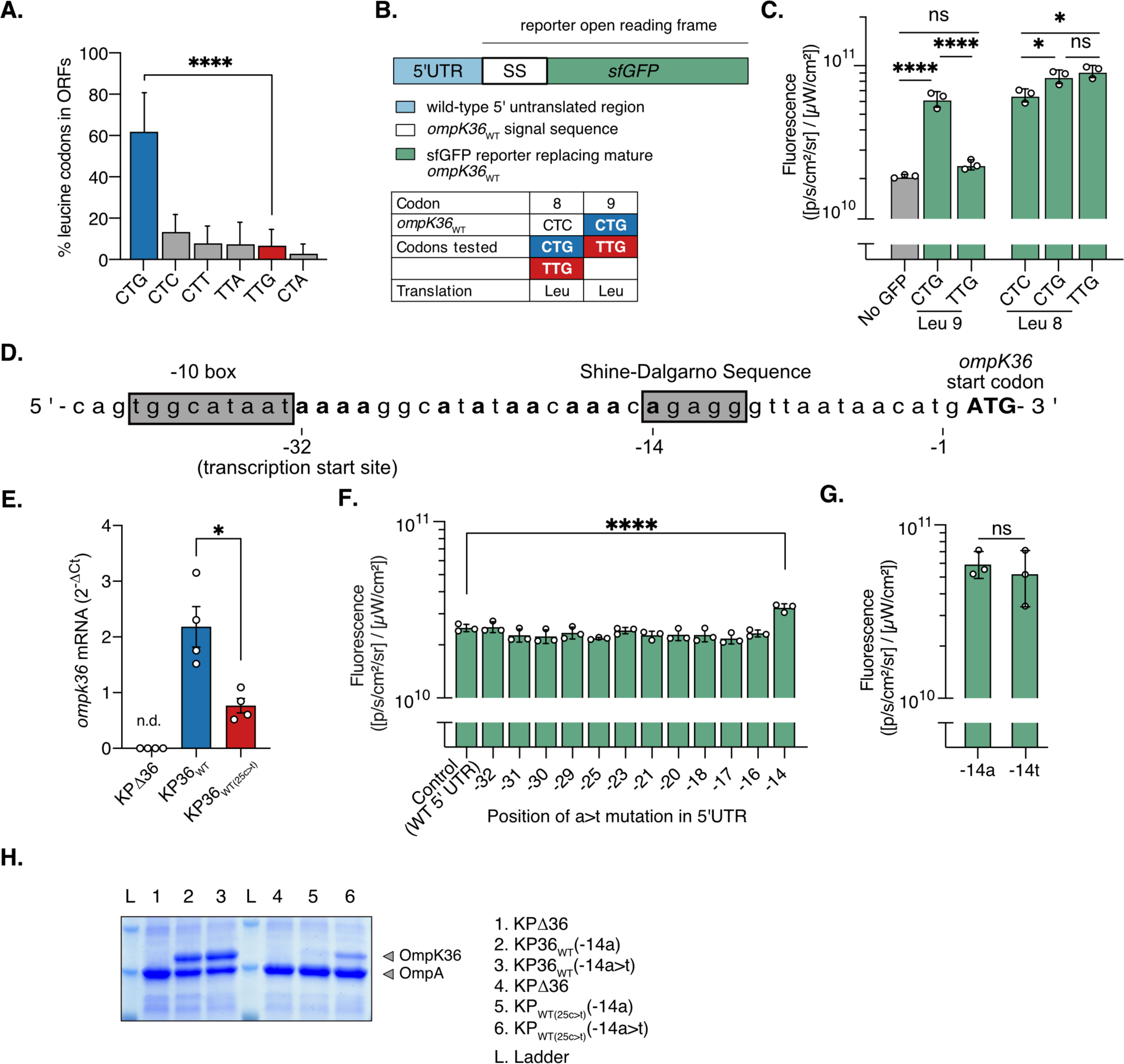
The 25c>t mutation results in a specific intramolecular interaction between -14a and 25u in the mRNA transcript. **A.** Proportion of leucine residues encoded by each of the six codons across individual open reading frames (ORFs) in the ICC8001 KP genome. A significant difference was observed in the use of CTG relative to TTG; significance was determined by paired T-test ****, p<0.0001. **B.** Schematic of the sfGFP reporter. The sequence encoding the mature OmpK36 protein was replaced with sfGFP, generating a chimeric fusion between the *ompK36*_WT_ signal sequence (Leu9 CTG) and sfGFP. The inset table describes the mutations that were introduced. **C.** Mutation of Leu9 CTG (*ompK36*_WT_) to TTG (*ompK36*_WT(25c>t)_) decreases expression of the sfGFP reporter. Mutations in Leu8 (CTC codon in *ompK36*_WT_) to CTG or TTG codons lead to similarly increased levels of reporter expression. Fluorescence was determined using IVIS SpectrumCT. Graphs show means of 3 biological repeats ± the standard deviation (SD). Statistical significance was determined by one-way ANOVA with Tukeýs multiple comparison test. ns, not significant; *, P<0.05; ****, P<0.0001. **D.** Sequence of the *ompK36* 5’UTR. Adenine bases in the 5’UTR (in bold) were individually mutated to thymine. **E.** RT-qPCR analysis of *ompK36* shows decreased transcript levels in the strain encoding *ompK36*_WT(25c>t)_ compared to *ompK36*_WT_. Bar charts show the mean ± the standard error of the mean (SEM) of 4 biological repeats. nd, not detected. ***P < 0.005 by paired Student’s *t* test. **F.** Expression of the sfGFP reporter encoding the *ompK36*_WT(25c>t)_ signal sequence is specifically increased upon mutation of the adenine in position -14 to thymine (-14a>t). The -14a marks the start of the Shine-Dalgarno sequence (SDS). Fluorescence was determined using IVIS SpectrumCT. Graphs show the means ± SD of 3 biological repeats. Statistical analysis was performed by one-way ANOVA with Dunnett’s multiple comparison post-test. ****, P<0.0001. **G.** Mutation of the adenine in position -14 to thymine (-14a>t) in the reporter construct encoding the *ompK36*_WT_ signal sequence (Figure 4B) does not impact sfGFP expression. Bar charts show the mean ± the standard deviation (SD) of 3 biological repeats. *ns*, not significant by paired Student’s *t* test. **H.** Representative Coomassie gel image of outer membrane preparations from isogenic KP strains with the indicated *ompK36* mutations in positions -14 (5’UTR) and 25 (ORF signal sequence). Mutation of the adenine in position -14 to thymine (-14a>t) restores expression of OmpK36 in the presence of the 25c>t mutation.

To investigate this, we generated a fluorescent ICC8001 reporter, in which the *ompK36* ORF was replaced with the *ompK36* signal sequence fused to sfGFP (Figure 4B). This reporter maintains the upstream promoter and regulatory regions at the monocistronic *ompK36* locus. Using sfGFP fluorescence as a proxy for protein expression, we found that the signal sequence containing the 25c>t mutation (TTG codon) significantly reduced expression compared to that with the WT sequence (CTG codon) (Figure 4C). This finding is consistent with the reduced OmpK36 abundance observed in the strain encoding *ompK36*_WT(25c>t)_ (Figure 2D). We then used the CTC leucine codon located at amino acid position eight (Leu8) to test the impact of codon usage on expression. We generated synonymous mutants where Leu8 was encoded by either of the alternative CTG (common) or TTG (rare) codons. Both Leu8 CTG and TTG codons resulted in similar sfGFP expression, which was significantly higher than with the CTC codon found in *ompK36*_WT_ (Figure 4C). This suggested that the underlying mechanism reducing OmpK36 expression via the 25c>t mutation is not related to codon usage limiting the efficiency of translation elongation.

### Reduced OmpK36 abundance is mediated by an interaction between the ORF and 5’UTR

Mutagenesis studies in *E. coli* have suggested that specific intramolecular RNA interactions between nucleotides in the 5’ end of an ORF and the upstream 5’UTR region can result in secondary structures that occlude the SDS, blocking ribosomal access (Kudla et al., 2009; Bhattacharyya et al., 2018). This disrupts the initiation of translation, reducing protein expression, and in turn results in mRNA degradation as transcription and translation are tightly coupled (Bhattacharyya et al., 2018). However, mutations reducing translation efficiency have not been identified as a naturally occurring mechanism tuning protein expression in adaptation to a host or environmental pressure. Nonetheless, we hypothesized that the reduced abundance of OmpK36WT(25c>t) may be due to an inhibitory mRNA secondary structure, mediated by a specific a/u base-pairing occurring between an adenine within the 5’ UTR of *ompK36* (Figure 4D) and the uracil encoded by 25t. We started by confirming the lower abundance of *ompK36* mRNA transcripts in the 25c>t mutant compared to WT by qRT-pCR (Figure 4E), a finding consistent with increased mRNA degradation. We then individually mutated each adenine to a thymine (starting from the end of the -10 box/transcription start site) in the 5’UTR of the sfGFP reporter containing the 25c>t mutation in the signal sequence. We hypothesized that the disruption of any base interaction between 25u and a position in the 5’UTR would be evidenced by increased sfGFP expression. No significant changes in fluorescence were observed in adenine substitutions up to and including the -16 position (Figure 4F). However, substitution of -14a, which marks the start of the SDS, abolished the effect of 25c>t on sfGFP signal suppression, with significantly increased fluorescence in the strain encoding the 14a>t mutation (p<0.001). In order to exclude the possibility that -14a>t had a non-specific effect on the functionality of the SDS due to enhanced ribosomal binding, we introduced -14a>t in the WT reporter (without 25c>t). Similar signal levels were observed in both reporters (-14a/25c and -14t/25c), suggesting that -14a>t alone does not affect protein expression (Figure 4G).

We next investigated the effect of the -14a>t mutation on OmpK36 abundance in the context of *ompK36*_WT_ and *ompK36*_WT(25c>t)_. As expected, no effect on abundance was observed when the mutation was introduced into *ompK36*_WT_ (Figure 4H). In contrast, introduction of the -14a>t mutation in the 5’UTR of *ompK36*_WT(25c>t)_ increased OmpK36 abundance (Figure 4H). These findings therefore suggest that the 25c>t mutation may result in an inhibitory mRNA secondary structure via a specific interaction with position - 14 in the 5’UTR.

### The 25c>t mutation induces an mRNA stem loop structure involving the Shine-Dalgarno sequence

To test the hypothesis that the 25c>t mutation results in an mRNA interaction between - 14a and 25u that obstructs the SDS and prevents ribosome recruitment, we probed the structures of the *ompK36*_WT_ and *ompK36*_WT(25c>t)_ mRNAs using dimethyl sulfate (DMS) mutational profiling with sequencing (DMS-MaPseq) (Zubradt et al., 2017). This technique uses DMS modifications of adenine and cytosine bases in the RNA, detected as mismatches during reverse transcription, to infer the accessibility of individual bases and subsequently build a model of the RNA structure. We found that adenine and cytosines within the SDS are more accessible to DMS in *ompK36*_WT_ mRNA than *ompK36*_WT(25c>t)_ mRNA both *in vitro* and *in vivo* (Figures 5A, 5B, Figure S1). Introducing the 24c>u mutation in addition to 25c>u reversed the DMS accessibility of the SDS to that resembling *ompK36*_WT_ (Figure 5C). As a negative control, we also tested a sample with no DMS treatment, which indeed showed minimal background signal (Figure 5D).

**Figure 5.**
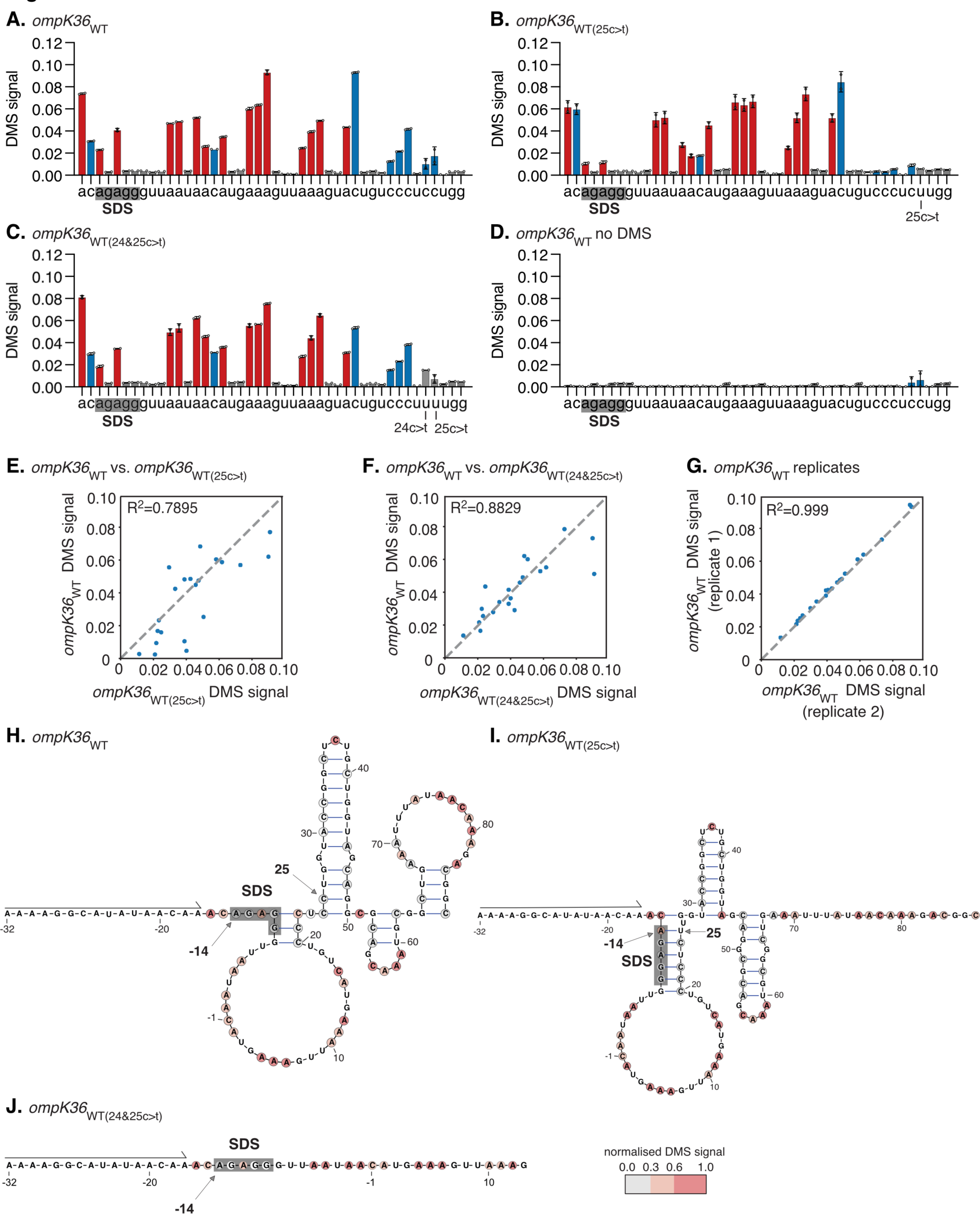
Position 25 in the *ompK36*_WT(25c>t)_ RNA induces a stem involving the Shine-Dalgarno sequence (SDS). **A-D** Dimethyl sulfate (DMS) signal per nucleotide from full length *in vitro-*transcribed and refolded *ompK36*, where higher values correspond to increased base accessibility. DMS signal and error bars for nt -16a through 28g are shown for *ompK36*_WT_ (**A**), *ompK36*_WT(25c>t)_ (**B**), *ompK36*_WT_(24&25c>t) (**C**) and DMS-untreated *ompK36*_WT_ (**D**). **E-G**. Comparison of DMS reactivities from nucleotides -16a through 28g between *ompK36*_WT_ and *ompK36*_WT(25c>t)_ (**E**), between *ompK36*_WT_ and *ompK36*_WT_(24&25c>t) (**F**), and between *ompK36*_WT_ replicates (**G**). Pearson correlations (R^2^) are shown. **H-J**. DMS-constrained structure models of the 5’ end of *ompK36*_WT_ (**H**), *ompK36*_WT(25c>t)_ (**I**) and *ompK36*_WT_(24&25c>t) (**J**). Nucleotides are colored by normalized DMS signal. The PCR primer binding sites where DMS information is unavailable are marked from positions -32 to -17. The SDS in RNA structures are highlighted in grey. Arrows indicate the -14 and +25 positions.

Correlation of the DMS signals between different pairs of *ompK36* variants (from positions -16 to +28) further demonstrated higher similarity between *ompK36*_WT_ and *ompK36*_WT_(24&25c>t) than between *ompK36*_WT_ and *ompK36*_WT(25c>t)_ (Figure 5E-G).

This implies that the RNA structure of the double mutant more closely resembles the WT. Indeed, DMS-driven structure models showed visually that the SDS is largely accessible for both the WT and the double mutant (although uncertainty exists in the model post position 13 of the double mutant) (Figures 5H-J, SI). In contrast, the SDS in *ompK36*_WT(25c>t)_ is sequestered into a stem structure, which incorporates the paired positions of -14 and 25. These results provide direct evidence that RNA structure occluding the SDS underlies the reduced expression of OmpK36 in strains encoding the 25c>t mutation. Altogether, these findings provide the precise molecular mechanism linking the occurrence of synonymous SNPs to altered OmpK36 abundance and meropenem resistance levels.

## Discussion

OmpK36 is a key porin in KP, reflected by its typically high abundance in the outer membrane. However, its role in facilitating antibiotic entry has exerted an evolutionary pressure favouring mutations that restrict this process (Vergalli et al., 2020). Using large clinical genome collections, we identified a single, recurrently-emerging synonymous base substitution at the 25^th^ nucleotide position of *ompK36* that increases carbapenem resistance by reducing OmpK36 expression. We observed that this 25c>t mutation usually occurs in the absence of L3 insertions, unless a 24c>t mutation which reverses the phenotype is also present, demonstrating that KP typically uses either OmpK36 pore constriction (via L3 insertions) or reduced OmpK36 abundance as a means for increasing resistance. Mechanistically, we show the significant impact a single synonymous substitution can have on protein expression, through the induction of a dramatic mRNA conformational change. To the best of our knowledge, this is the first time an adaptive mutation that restricts translation initiation has been shown to occur naturally.

In unravelling the molecular mechanism, our *in vitro* mutagenesis experiment showed that reduced OmpK36 expression via the 25c>t mutation is the result of a specific interaction between 25u in the mRNA and the first adenine base at the upstream SDS (-14a). The interaction was confirmed by solving mRNA structures with DMS-MaPseq. This technique revealed large conformational differences in the structures of the *ompK36*_WT_ and *ompK36*_WT(25c>t)_ transcripts, including the presence of a stem loop structure sequestering the SDS only in the *ompK36*_WT(25c>t)_. We propose that this stem loop structure, triggered by the interaction between 25u and -14a, impedes ribosomal binding and thereby reduces translation efficiency. Further evidence for this proposed mechanism also came from the mRNA transcript model of *ompK36*_WT_(24&25c>t) (these double mutations also having been observed among clinical isolates), which again had a different conformation but showed no stem loop structure obscuring the SDS. This could explain the reversal of the phenotypes observed in this mutant compared with that encoding only 25c>t. Indeed, previous studies have demonstrated that experimentally-induced mRNA secondary structures near the SDS decrease protein levels using a synthetic library of synonymous GFP variants (Kudla et al., 2009) and, more recently, using two endogenous genes in *E. coli* (Bhattacharyya et al., 2018). Crucially, however, it has been proposed that the base composition at the start of genes has evolved to minimise the formation of these structures, as evidenced by a genome-wide analysis in *E. coli* (Bhattacharyya et al., 2018). This appears to be the case in the *ompK36*_WT_ transcript, in which the SDS lies in a largely open conformation. However, the mechanism behind the increased resistance of the 25c>t mutant (and reversal of this phenotype in the 24&25c>t mutant) extends our understanding of this system by demonstrating that mutations which induce alternative secondary structures can also be selected during adaptation. Importantly it also provides a demonstration of the functional use of these inhibitory mRNA structures in regulation among naturally occurring and clinically relevant populations.

In keeping with the key role of OmpK36 in maintaining KP physiology, we demonstrated that the 25c>t mutation decreased bacterial replication in a murine pneumonia model. This correlated with reduced levels of serum IFNγ which plays an important role in KP lung infection in mice (Yoshida et al. 2001; Moore et al., 2002). Our phylogenetic analyses suggested that this level of attenuation may be enough to impact on transmission, with most isolates possessing this mutation forming singletons or pairs in the ST258/512 phylogeny (i.e. not being part of larger clonal expansions). Moreover, despite its frequent emergence, the total prevalence of the 25c>t mutation among the ST258/512 genome collection remained at 1.2%. This is similar to the prevalence of loss-of-function mutations in *ompK36* (0.7%) but is in sharp contrast to L3 insertions, which have spread widely via large clonal expansions to reach a prevalence of 47.0%. Nevertheless, despite the limited clonal expansion of 25c>t, our identification of this mutation in clinical isolates demonstrates that it does not preclude infection in patients.

Using our translational model of pneumonia, we could further show that the increased resistance conferred by the 25c>t mutation was enough to result in a clinical impact, thereby explaining its recurrent emergence (despite the fitness cost). Mice infected with KP expressing *ompK36*_WT_+KPC-2 could be successfully treated with meropenem, while those infected with KP possessing *ompK36*_WT(25c>t)_+KPC-2 could not. We therefore propose that the repeated emergence of 25c>t is driven by antibiotics, possibly due to prolonged exposure and/or subtherapeutic dosing; both are commonly encountered among critically ill patients, in whom antibiotic pharmacokinetics are difficult to predict at an individual patient level (Roberts et al., 2014). Indeed, porin modifications occurring within the time course of a single infection have been reported previously (Elliott et al., 2006; Yoshino et al., 2021), demonstrating potential for their emergence and selection given strong antibiotic pressure. Increased host susceptibility due to co-morbidity and impaired immune responses in the population most at risk of KP disease could also render the associated fitness cost of 25c>t less significant.

While the frequency of 25c>t has remained low, we propose that its ongoing *de novo* emergence, together with the spectrum of other known OmpK36 modifications, impose a significant impact on patient treatment. Moreover, porin mutations that reduce carbapenem entry are not restricted to KP and are found in other WHO critical priority organisms such as *Pseudomonas aeruginosa* and *Escherichia coli* (Bajaj et al., 2016; Lister et al., 2009). This highlights the need for the development of novel effective antibiotic therapies that circumvent reliance on diffusion through porins. While this has been achieved via the newly licensed cephalosporin, cefiderocol, which instead enters via siderophore uptake systems (Kohira et al., 2016), additional drugs are still needed.

In summary, our combined genomic, experimental and translational approaches have uncovered a novel mechanism underpinning carbapenem resistance, mediated by synonymous mutations that alter *ompK36* mRNA secondary structure. The associated dynamics of emergence and expansion of these mutations are at the heart of an evolutionary conflict balancing resistance and virulence requirements in KP. We propose a central role for combining genomic surveillance with *in vitro* and translational evaluation to further our understanding of resistance mutations, and to design and target our clinical interventions accordingly.

## Acknowledgments

We would like to thank Michelle Yeap for making the vector to delete *ompK35,* Izabela Glegola-Madjeska for support with animal studies and the Pathogen Informatics group from the Wellcome Sanger Institute for informatics support. We also thank Edward Feil and members of the JPI-AMR-funded SPARK consortium (grant MR/R00241X/1) for access to genomes and metadata used in this study.

## Author contributions

JLCW and SD conceived the study. SD performed bioinformatics and genomic analyses. JLCW performed the molecular biology and biochemistry experiments. WWL performed the outer membrane preparations. JLCW, JSG and WWL performed the animal experiments under the supervision of GF. JSG performed cytokine measurements. FM, TG and GMR performed the meropenem resistance assays. JZW performed the DMS-MaPseq under the supervision of SR. Data analysis was carried out by JLCW, SD and JSG. JLCW, SD and GF wrote the manuscript. All authors reviewed and edited the manuscript (JLCW, SD, JSG, JZW, WWL, FM, TG, GMR, SJB, AC, DMA, SR and GF).

## Supplementary Figure Legends

**Figure S1. Position 25 in the *ompK36*_WT(25c>t)_ RNA forms a stem with the Shine-Dalgarno sequence (SDS) *in vivo*.**

**A.** Dimethyl sulfate (DMS)-constrained predicted structures of *in vivo ompK36*_WT_ (top) and *ompK36*_WT(25c>t)_ (bottom) RNA molecules. Nucleotides are colored by normalized DMS-MaPseq mutation rate (DMS signal), where higher values correspond to increased base accessibility. The PCR primer binding site where DMS information is unavailable is highlighted. *In vivo* RNA structures correspond with *in vitro* transcribed RNA structures (Figure 5A&B) with SDS sequestration in the *ompK36*_WT(25c>t)_ but not *ompK36*_WT_ RNA molecule.

**B.** Correlation of DMS signals for each nucleotide between two biological replicates for *in vivo ompK36*_WT_ (top) and *ompK36*_WT(25c>t)_ (bottom).

**C.** DMS signal per nucleotide of *in vivo ompK36*_WT_ (top) and *ompK36*_WT(25c>t)_ (middle) and DMS-untreated *ompK36*_WT_ (bottom) RNA molecules. The SDS are highlighted in grey boxes.

## Supplementary Table Legends

**Table S1.** Metadata and genotyping data for a curated collection of 1450 KP ST258/512 genomes.

**Table S2.** Metadata and genotyping data for 16,086 KP genomes available in Pathogenwatch.

## Methods

### Genome collections

We used two genome collections to characterise the diversity and phylogenetic distribution of *ompK36* (and *ompK35*) variants. The first comprises 1450 public KP genomes belonging to STs 258, 512 and other nested STs, together with curated metadata obtained from associated publications (**Table S1**). We obtained raw sequence reads if available (1340 isolates), and short-read assemblies from the remainder (110 isolates). Assemblies were generated for all isolates with available raw sequence data using SPAdes v3.9.0 (Bankevich et al., 2012), and annotated with Prokka v1.14.5 (Seemann, 2014). Kleborate v1.0.0 (Lam et al., 2021) was used for confirming the ST of each genome and determining the resistance gene content.

The second collection comprises a public genome collection of 16,086 KP available in Pathogenwatch together with available metadata and genotyping data (Argimon et al., 2021) (https://pathogen.watch/genomes/all?genusId=570&speciesId=573) (**Table S2**).

### Phylogenetic analysis of the ST258/512 collection

To generate a phylogenetic tree of the ST258/512 collection, we first used the “to_perfect_reads” function within Fastaq v3.17.0 (https://github.com/sanger-pathogens/Fastaq) to generate pseudo sequence reads for isolates where only an assembly was available. We then mapped all sequence reads to the reference genome, NJST258_1 (accession CP006923) (Deleo et al., 2014), using Burrows Wheeler Aligner v0.7.17 (Li & Durbin, 2009). A pipeline comprising SAMtools mpileup v0.1.19 (Li et al., 2009) and BCFtools v0.1.19 was used to call SNPs and generate a pseudo-genome alignment. Gubbins v2.4.1 (Croucher et al., 2015) was used to remove recombined regions from the alignment and generate a maximum likelihood tree with the remaining variable positions. The phylogenetic tree was rooted using an outgroup isolate from a closely-related ST, ST895 (accession SRR5385992), which was later removed from the tree.

### Identification and characterisation of *ompK35 and ompK36* genes

The *ompK35 and ompK36* genes were identified in all short-read assemblies by performing BLASTn v2.6.0 (Altschul et al., 1990) with a query gene from the reference genome, ATCC43816 (the parental strain of ICC8001 (Wong et al., 2019)) (accession CP009208). We required a single hit of each gene per genome that matched ≥10% of the query length, possessed ≥90% nucleotide similarity and contained a start codon in order to unambiguously identify the gene. Nucleotide sequences were translated to protein sequences in Seaview v4.7 (Gouy et al., 2010) using the standard genetic code, and the protein lengths were determined. Protein sequences were predicted to be intact if they contained a signal sequence, a L3 sequence and sixteen beta-barrel sequences, as determined using BLASTx v2.6.0 (Gish & States, 1993). Intact protein sequences of each porin were aligned using MUSCLE v3.8 (Edgar, 2004) and the different variants present were identified, taking into account all amino acid substitutions, insertions and deletions.

### Analysis of *ompK36* variation

The alignment of intact *ompK36* nucleotide sequences from the ST258/512 collection was curated manually to ensure accurate positioning of bases around the L3 region (some of which were misaligned due to indels). We then used the sequences from the curated alignment to infer the likely base harboured by each internal node of the ST258/512 phylogenetic tree at each position in *ompK36*. This ancestral reconstruction was performed with PastML v1.9.30 (Ishikawa et al., 2019) using maximum parsimony with the accelerated transformation (ACCTRAN) model. Using the predicted states of all internal nodes and the known bases in all terminal nodes, we determined the total number of changes at each position that had occurred across the ST258/512 tree.

### Codon usage

The frequency of different leucine codons was determined across all protein-coding genes in the ATCC43816 reference genome, as well as all annotated assemblies in the ST258/512 collection, using the EMBOSS v6.6.0.0 “cusp” tool (http://emboss.sourceforge.net/apps/release/6.3/emboss/apps/cusp.html).

### 5’ untranslated region analysis

The 500bp upstream region of *ompK36* was analysed and annotated in Fig. 4D using BPROM (Solovyez & Salamov, 2011). The SDS was identified manually by consensus with other KP genes.

### General cloning, molecular biology and strain generation

All *in vitro* assays and animal infections with KP were carried out using ICC8001, a strain derived by animal passage from ATCC43816 (Wong et al., 2019).

A homologous recombination technique resulting in scarless markerless mutants was used to generate all strains based on the Standard European Vector Architecture platform pSEVA612S (JX560380.2) mutagenesis plasmid. Briefly, mutants were generated in two sequential recombination steps. The first step integrates the mutagenesis plasmid into the genome generating a merodiploid. The mutagenic plasmid was introduced by three part conjugation and homologous recombination promoted by lambda-recombinase expressed from a helper plasmid. The second step induces double strand DNA breaks by induction (L-arabinose) of I-SceI expression from the helper plasmid. Double stranded breaks are induced at I-SceI sites located on the chromosomally integrated mutagenesis vector, flanking the final integration region. This technique has been used to efficiently recombineer in KP (Wong et al., 2019) and other *Enterobacteriaceae* (Berger et al., 2017) as described in respective manuscripts.

Mutagenesis vectors were generated by routine molecular biological techniques. These included plasmid preparation (Monarch Plasmid Miniprep Kit, NEB), genomic DNA extraction from bacterial cells (DNEasy, QIAGEN), polymerase chain reaction (Q5 High-Fidelity 2x Master Mix and OneTaq, Quick-Load 2X Master Mix, both NEB), PCR clean-up (Monarch PCR and DNA Cleanup Kit, NEB) and gel extraction (Monarch Gel Extraction Kit, NEB), site directed mutagenesis (KLD enzyme blend, NEB) and Gibson assembly (Gibson Assembly Master Mix, NEB). Sanger sequencing was carried out to check vector construction and seamless integration into the genome (Eurofins Genomics). Custom primers in this study were made by Sigma-Aldrich.

### *ompK35* mutagenesis

*ompK35* wild-type ORF (+500bp flanking regions) was cloned from ICC8001 genomic DNA with primers (P) 1/P2 and ligated by Gibson Assembly into pSEVA612S, linearised with P3/P4. The ORF was removed by inverse PCR to generate the genomic deletion vector (P5/P6) and plasmid recircularized from PCR product (KLD enzyme blend, NEB).

### *ompK36* mutagenesis

ICC8001Δ*ompK36* was generated in previous work (Vector 5 in Wong et al., 2019). *ompK36* wild-type ORF (+500bp flanking regions) was cloned from ICC8001 genomic DNA with primers P7/P8 and ligated by Gibson assembly into pSEVA612S, linearised with P3/P4. In screening colonies by Sanger Sequencing we found the 25c>t mutation and this was not generated by site directed mutagenesis.

### sfGFP vectors and -14a>t vectors

#### Leucine 9 CTG-sfGFP (wild-type) and Leucine 9 TTG-sfGFP (25c>t transition)

The wild-type and 25c>t Sec signal sequences were cloned from the *ompK36* vectors generated above with P9/P10. sfGFP was amplified from our previously published glmS site insertion vector (Vector 13 in Wong et al., 2019) with P11/P12 and chimeric fusion generated in the mutagenesis vector ligated by Gibson Assembly.

Leucine 8 CTG-sfGFP and Leucine 8 TTG-sfGFP were generated by site-directed mutagenesis using P13/P14 and P15/P16 respectively using the Leucine 9 CTG-sfGFP (wild-type) as the template.

#### 5’ untranslated region mutagenesis of Leucine 9 TTG-sfGFP (25c>t transition)

Mutants were generated by site-directed mutagenesis as follows: -32(a>t) (P17/P18), -31(a>t) (P17/P19), -30(a>t) (P17/P20), -29(a>t) (P17/P21), -25(a>t) (P22/P23), -23(a>t) (P22/P24), -21(a>t) (P22/P25), -20(a>t) (P22/P26), -18(a>t) (P22/P27), -17(a>t) (P22/P28), -16(a>t) (P22/P29), -14 a>t(a>t) (P22/P30) using leucine 9 TTG sfGFP as the template. The same primer pair (P22/P30) was used to introduce the -14a>t mutation into Leucine 8 CTG-sfGFP, *ompK36WT* and *ompK36WT(25c>t)* vectors.

### Introduction of pKpQIL KPC-2 plasmid

We introduced pKpQIL KPC-2 by conjugation using an *E. coli* donor following our previously published protocol (Wong et al., 2019).

### Meropenem Minimum Inhibitory Concentrations (MIC)

The meropenem MIC of strains was assessed by broth microdilution by broth microdilution in biological triplicate according to the ISO 20776-1:2019 standard and the median value taken as the final MIC.

### Outer membrane preparations and gel electrophoresis

Outer membrane proteins were purified according to a previously described protocol with several modifications (Wise et al., 2018). Saturated overnight cultures of bacteria were grown in LB (10g/L NaCl) and sonification was performed at 25% amplitude for 10 bursts of 15 seconds each. 10 μg of protein was separated by SDS-PAGE using 12% acrylamide Mini-protean TGX precast gels (Bio-Rad, USA). Gels were stained with Coomassie (Sigma-Aldrich) and imaged on a ChemiDoc XRS+ (Biorad, USA).

### Animal work

8-10 week old, female, 18-20g BALB/c mice were purchased from Charles River, UK. All animal work took place in the institution animal facility (Association for Assessment and Accreditation of Laboratory Animal Care accredited) under the auspices of the Animals (Scientific Procedures) Act 1986 (PP7392693). Work was approved locally by the institutional ethics committee.

### Housing and Randomisation

Upon arrival animals were independently randomized into cages of 5 animals and housed for an acclimatisation period of 1 week. Mice received food and water *ad libitum* and were housed in a 12hour/12hour light dark cycle. Identification of animals within groups was achieved by ear notching without anaesthesia at least 24 hours in advance of procedures.

### Intratracheal administration of inoculum and infection

Anaesthesia was induced by intraperitoneal injection (BD Microlance 27G 13mm needle) of 80mg/kg ketamine and 0.8mg/kg medetomidine. Pre-intubation body temperature was maintained by contact with a heat mat (Harvard Apparatus, U.K) and eye lubricant (2.0mg/g carbomer, Alcon UK) was applied.

Intubation was achieved as previously described (Wong et al., 2019) by placement of a 21G catheter (21G IV peripheral catheter (Insyte) BD Medical) using a fibreoptic intubation set (Kent Scientific) following the enclosed instructions.

Inoculum (dose defined in graphs) was prepared by dilution of saturated overnight cultures into phosphate buffered saline (PBS) (total inoculum volume 50ul) and following inspiration it was dispersed with 2 x 200ul air flushes. An aliquot of inoculum was enumerated for each administration and all inoculums were ±10% of the stated figure. Mice recovered at 32°C (Warm Air System, Safety Cabinet Version red, VetTech, UK) until spontaneous movement and received 0.8mg/kg subcutaneous (BD Microlance 27G 13mm needle) atipamezole to reverse the ɑ-agonist into the neck scruff.

At the experimental end-point animals were anesthetized with ketamine 100mg/kg and 1mg/kg medetomidine by intraperitoneal injection (BD Microlance 27G 13mm needles). Following the induction of anaesthesia, blood was collected by a transdiaphragmatic inferior approach cardiac puncture (BD Microlance 25G 25mm needles) and animals were then humanely killed, under anaesthesia, by cervical dislocation.

An aliquot of blood (20ul) was taken and diluted into 180ul of hypotonic lysis solution (1mM EDTA in water) for enumeration of bacterial counts. The rest was placed into a microtainer (BD Microtainer for serum collection, BD Medical) and allowed to clot for 1 hour. This was then spun according to the manufacturer’s guidance and aliquoted into cryobanking tubes (Greiner Bio-One) and frozen at -80℃ until analysis.

Lungs were dissected out, weighed and then homogenised in gentlemacs C-tubes (Miltenyi Biotec) containing 3 ml complete RPMI media (RPMI supplemented with 10% heat inactivated FBS, 10 mM HEPES and 1mM Sodium Pyruvate) using a gentleMac Octo dissociator (Miltenyi Biotec) using the program m_lung_1 twice.

Bacteria were enumerated from the lung homogenate and blood by dilution in PBS and plating onto solidified LB agar containing 50ug/ml rifampicin (Merck, UK).

### Antibiotic delivery

We combined the meropenem (100mg/kg) with cilastatin (100mg/kg) (Merck, UK), a renal dihydropeptidase inhibitor, to reduce the *in vivo* metabolism of meropenem as the murine enzyme isoform has increased activity against this carbapenem compared to human enzyme. Drugs were diluted in dH2O. dH2O alone was administered to vehicle control animals. Doses were delivered at 6 hourly intervals as previously employed in a murine carbapenemase producing KP model (Ota et al., 2020). The drug mixture was delivered by intraperitoneal infection (BD Microlance 27G 13mm needles) to animals at 6 hourly intervals and the iliac fossa used was alternated between doses.

### Cytokine bead assay

Serum IFN-γ was assessed using a custom-made mouse panel 13-plex kit (LEGENDplex, BioLegend) following the manufacturer’s instructions. Cytokine levels were acquired using a FACSCalibur flow cytometer (BD Biosciences), and analyses were performed using LEGENDplex data analysis software (BioLegend). A biological repeat was considered valid only if at least 80% of all the samples run were above the limit of detection; any values below the detection limit within a repeat such as this were assumed to be the lowest value detectable by the assay for statistical analysis. Values below the detection limit in graphs were displayed as 1 for visualisation.

### RNA isolation and reverse transcription quantitative PCR (RT-qPCR)

300 µl of an LB overnight culture of KP were treated with RNAprotect Bacteria reagent (Qiagen) and centrifuged at 5000 xg for 10 min. The bacterial pellet was digested in 100 µl TE buffer with 15 mg/ml Lysozyme (Sigma-Aldrich) and 20 µl Proteinase K (Qiagen) for 10 min according to manufacturer’s guidelines. RNA was isolated using the RNeasy minikit (Qiagen) following the manufacturer’s instructions, and RNA concentration was determined using a Nanodrop 2000 spectrophotometer (ThermoFisher Scientific). 1 µg of RNA was treated with RNase-free DNase (Sigma-Aldrich) for 1 h at 37 °C and cDNA was then synthesized using a Moloney murine leukemia virus (MMLV) reverse transcription kit with random primers following the manufactureŕs protocol (Promega). To check for the presence of remaining DNA, a reaction mixture without the MMLV reverse transcriptase was also included (NRT). qPCR was performed using the Power Up SYBR Green master mix (ThermoFisher Scientific) and the following primers 31/32 (*ompK36*) and 33/34 (*rpoB*). The assay was run on a StepOnePlus System (Applied Biosystems) and results were analysed using the StepOne software (Applied Biosystems). Relative gene expression levels were analyzed by using the 2^’(^*^CT^*(where *CT* is threshold cycle) method.

### IVIS sfGFP fluorescence measurement

Saturated overnight cultures of strains were diluted 1:2000 in PBS. 20ul of suspension was plated onto an agar plate and allowed to dry. Following overnight incubation (16 hours exactly) plates were removed and imaged on an IVIS Spectrum CT imaging platform (1 second exposure, Ex 465nm/Em520nm). Fluorescence over the area of the spot was quantified using Living Image Software v5.4.3 (Perkin Elmer).

### Determination of RNA structures using DMS-MaPseq

RNA structures of *ompK36*_WT_, *ompK36*_WT(25c>t)_ and *ompK36*_WT_(24&25c>t) were determined using DMS-MaPseq similarly to as described previously (Zubradt et al., 2017). The principle of this method is that DMS adds methyl groups to the Watson-Crick faces of unpaired adenines and cytosines in the RNA. These methyl adducts are detected during reverse transcription using the TGIRT-III reverse transcriptase, which incorporates random mutations in the complementary cDNA at the corresponding nucleotides. These cDNA molecules are then amplified by PCR, followed by massively parallel sequencing. Mutations observed on a single DNA molecule correspond to the DMS-reactive bases on the parent RNA molecule if background levels of DMS-MaPseq can be observed to be negligible in untreated samples. *In vitro* (*ompK36*_WT_, *ompK36*_WT(25c>t)_ *and ompK36*_WT_(24&25c>t)) and *in vivo* (*ompK36*_WT_ and *ompK36*_WT(25c>t)_) DMS modification were performed as described below to solve the RNA structures.

### In vitro DMS modification

Plasmids encoding the wildtype and *ompK36* mutants were isolated using miniprep (Zymo) and amplified by PCR using a forward primer containing the T7 promoter sequence (TAATACGACTCACTATAGGAAAAGGCATATAACAAACAGAGGG) and the reverse primer (AAGAGTATACCAGCGAGGTTAAACCGGAC). The PCR product was used as a T7 Megascript *in vitro* transcription (ThermoFisher Scientific) template according to manufacturer’s instructions. Next, the RNA was purified using RNA Clean and Concentrator^TM^-5 (Zymo). 10µg RNA was denatured at 95°C for 1 min. Denatured RNA was refolded by incubating in 340mM sodium cacodylate buffer (Electron Microscopy Sciences) and 5mM MgCl^2+^, such that the volume was 97.5µl, for 20 min at 37°C. Then, 2.5% DMS (Millipore-Sigma) was added and incubated for 4 mins at 37°C with whole shaking at 800 r.p.m. on a thermomixer. Subsequently, DMS was neutralized by adding 60µl β-mercaptoethanol (BME) (Millipore-Sigma). The DMS-modified RNA was purified using RNA Clean and Concentrator^TM^-5 (Zymo) and eluted in 10µl water.

#### In vivo DMS modification

500µl of exponentially growing *E. coli* were incubated with 10µl DMS for 3 mins at 37°C shaking at 800 r.p.m. on a thermomixer. DMS was quenched by adding 500µl 30% BME, followed by a 3 min 30% BME wash and 1x PBS wash. Then, bacterial pellets were resuspended and incubated at room temperature for 5 mins in 500µl RNAprotect® bacteria reagent (QIAGEN). Samples were centrifuged at 16000g for 5 mins and supernatant removed. Pellets were resuspended in 100µl of 15mg/ml lysozyme solution plus 20µl proteinase K and mixed by vortex for 10 seconds. Samples were incubated at room temperature for 10 mins, vortexing every 2 mins for 10 seconds. Next, samples were added with 350µl of buffer RLT plus BME (10µl BME for every 1ml of RLT), vortex to mix. Then, 250µl of 96-100% ethanol was added to samples. RNA was extracted using the RNeasy® Mini kit (QIAGEN) following manufacturer’s instructions. DNA was digested from 5-10µg of total RNA per sample using the TURBO DNA-free kit (ThermoFisher Scientific) and purified using RNA Clean and Concentrator^TM^-5 kit. Following that, ribosomal RNAs were depleted using the Ribominus^TM^ Transcriptome Isolation kit for bacteria (ThermoFisher Scientific) following manufacturer’s instructions. RNA was purified using RNA Clean and Concentrator^TM^-5 (Zymo) and eluted in 10µl water.

#### RT-PCR and sequencing of DMS-modified RNA

To reverse transcribe, rRNA-depleted total RNA or *in vitro*-transcribed RNA purified from the previous steps was added to 4µl 5x FS buffer, 1µl dNTP, 1µl 0.1M DTT, 1µl RNase Out, 1µl 10uM reverse primer (CCTGAACGTTGTATTCCC) and 1µl TGIRT-III (Ingex). The reaction was incubated for 1.5h at 60°C. Then, to degrade the RNA, 1ul 4M NaOH was added and incubated for 3min at 95°C. The cDNA was purified in 10µl water using the Oligo Clean and Concentrator^TM^ kit (Zymo). Next, 1µl of cDNA was amplified using Advantage HF 2 DNA polymerase (Takara) for 30 cycles according to the manufacturer’s instructions (forward primer: GGAAAAGGCATATAAC; reverse primer: CCTGAACGTTGTATTCCC). The PCR product was purified using E-Gel^TM^ SizeSelect^TM^ II 2% agarose gel (Invitrogen). RNA-seq library for 300bp insert size was constructed following the manufacturer’s instructions (NEBNext Ultra^TM^ II DNA Library Prep Kit). The library was loaded on an iSeq-100 Sequencing flow cell with iSeq-100 High-throughput sequencing kit and the library was run on iSeq-100 (paired-end run, 151 x 151 cycles).

### Data analysis and visualisation

Metadata, including *ompK36* variation, were mapped onto a phylogenetic tree of the ST258/512 collection using iTOL v5.7 (Letunic & Bork, 2021) for use in figures. An interactive visualisation of this genome collection can also be accessed using Microreact v157 (Argimon et al., 2016): https://microreact.org/project/1vWbaqARPRNc55n4yfdLyQ-ompk36.

Graphs and statistical analysis was carried out in GraphPad Prism v9.0.0 for Mac (GraphPad Software, San Diego, California USA, www.graphpad.com). Data were analysed for normal distribution (based on D’Agostino-Pearson or Shapiro-Wilk normality tests) and if not normally distributed a base 10 logarithmic transformation was applied before statistical analysis. When normality was not achieved by transformation, a non-parametric test was applied. Statistical tests applied for each analysis are described in associated figure legends.

Custom images of mice in Fig. 3 were made with Biorender (created with <u>BioRender.com</u>). Images were edited and compiled into figures in Affinity Designer v1.8.4. (Serif Europe Ltd).

### Primers

P1 GCCAGTATAGGGATAACAGGGTAATCCGGTACCGCCCGGTGTCTG P2 CGCCGGATTACCCTGTTATCCCTAGGAAATCACAATTATGTTACG P3 ATTACCCTGTTATCCCTATAC

P4 TAGGGATAACAGGGTAATCCG P5 TCTGCAGTACACCCCTTC

P6 CATTATTTATTACCCTC

P7 GATTACGCCGGATTACCCTGTTATCCCTAGGCCTAATTGATTGATTAATAG P8 GGCCAGTATAGGGATAACAGGGTAATAGCCCCACAGGTTGACCAGC

P9 GTTGCAAGCTGCATAAC

P10 CGCATTTGCTGCGCCTGCTAC

P11 GTAGCAGGCGCAGCAAATGCGCGTAAAGGCGAAGAGCTG P12 GTTATGCAGCTTGCAACCGCGAAGTAATCTTTTCG

P13 TTAACTTTCATGTTATTAACCCTCTGTTTGTTATATG P14 AGTACTGTCCCTGCTGGTACCGG

P15 AGTACTGTCCTTGCTGGTACCGG P16 TTAACTTTCATGTTATTAACCCTCTG

P17 GCATATAACAAACAGAGGGTTAATAAC P18 CTTTAATTATGCCACTGC

P19 CTTATATTATGCCACTGC P20 CTATTATTATGCCACTGC P21 CATTTATTATGCCACTGC P22 CTTTTATTATGCCACTGC

P23 GCTTATAACAAACAGAGGGTTAATAAC

P24 GCATTTAACAAACAGAGGGTTAATAAC P25 GCATATTACAAACAGAGGGTTAATAAC P26 GCATATATCAAACAGAGGGTTAATAAC P27 GCATATAACTAACAGAGGGTTAATAAC P28 GCATATAACATACAGAGGGTTAATAAC P29 GCATATAACAATCAGAGGGTTAATAAC P30 GCATATAACAAACTGAGGGTTAATAAC P31 CCAGACCTACAACGCAACT

P32 CGCTCCAGATCCTTACCTTTAG P33 AAGGCGAATCCAGCTTGTTCAGC

P34 TGACGTTGCATGTTCGCACCCATCA

## Data availability

Raw sequence data or assemblies from the extended ST258/512 collection were published in multiple manuscripts and obtained from the European Nucleotide Archive (ENA). Genome assemblies from the larger KP collection were obtained from Pathogenwatch (https://pathogen.watch/genomes/all?genusId=570&speciesId=573). A full list of accession numbers for all data is available in Tables S1 and S2.

