## Supplementary figures and images for "Recurrent emergence of carbapenem resistance in *Klebsiella pneumoniae* mediated by an inhibitory *ompK36* mRNA secondary structure"

### Supplementary Figure 1

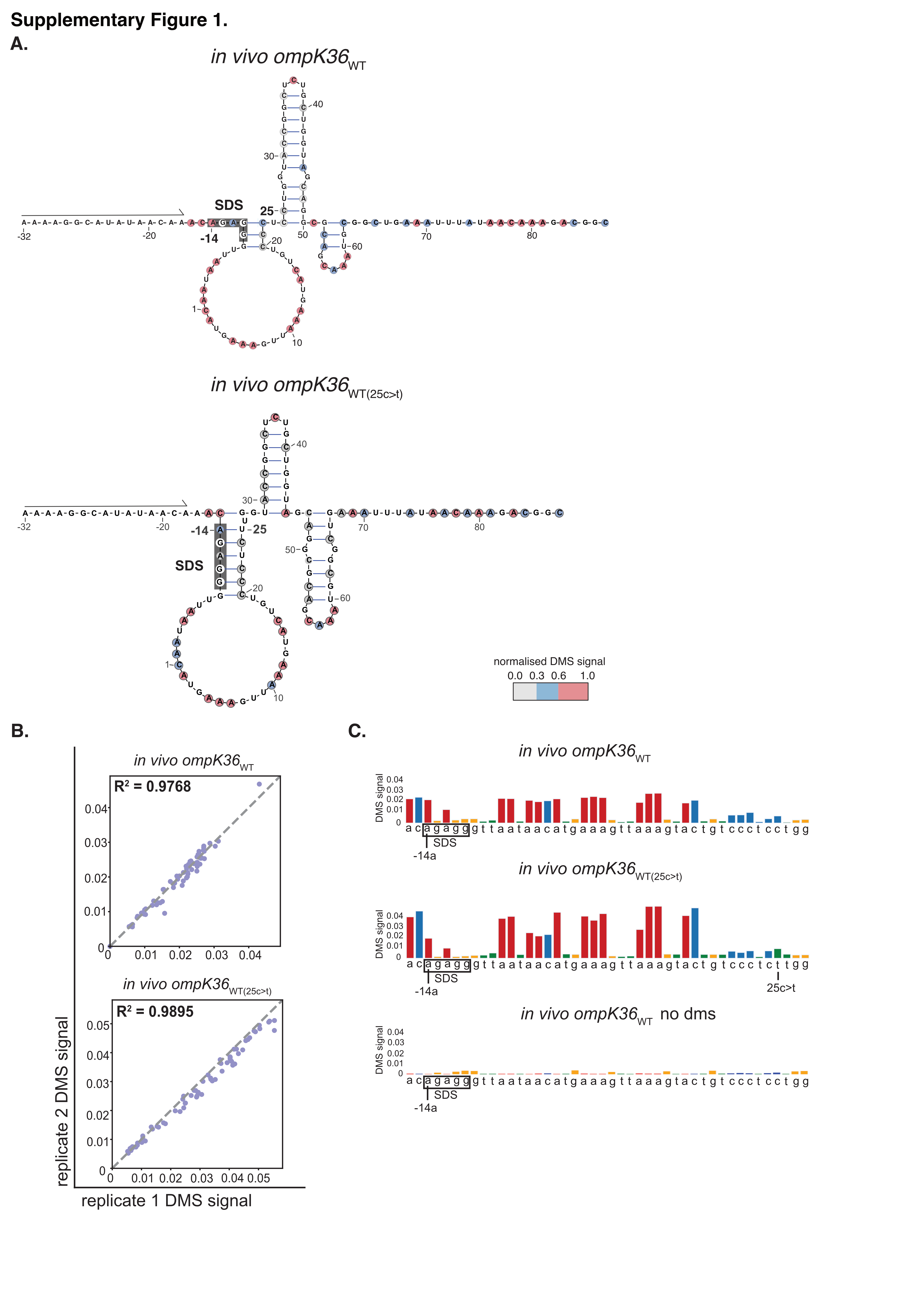
